# The Use of a SOX10 Reporter Towards Ameliorating Oligodendrocyte Lineage Differentiation from Human Induced Pluripotent Stem Cells

**DOI:** 10.1101/2023.12.01.569591

**Authors:** Valerio E.C. Piscopo, Alexandra Chapleau, Gabriela J. Blaszczyk, Julien Sirois, Zhipeng You, Vincent Soubannier, Geneviève Bernard, Jack P. Antel, Thomas M. Durcan

## Abstract

Oligodendrocytes (OLs) are key players in the central nervous system, critical for the formation and maintenance of the myelin sheaths insulating axons, ensuring efficient neuronal communication. In the last decade, the use of human induced pluripotent stem cells (iPSCs) has become essential for recapitulating and understanding the differentiation and role of OLs *in vitro*. Current methods include overexpression of transcription factors for rapid OL generation, neglecting the complexity of OL lineage development. Alternatively, growth factor-based protocols offer physiological relevance but struggle with efficiency and cell heterogeneity. To address these issues, we created a novel SOX10-P2A-mOrange iPSC reporter line to track and purify oligodendrocyte precursor cells (OPCs). Using this reporter cell line, we analyzed an existing differentiation protocol and shed light on the origin of glial cell heterogeneity. Additionally, we have modified the differentiation protocol, towards enhancing reproducibility, efficiency, and terminal maturity. Our approach not only advances OL biology but holds promise to accelerate research and translational work with iPSC-derived OLs.

**Main Points:**

- The differentiation of iPSCs in Oligodendrocyte Precursor Cells (OPCs) and Oligodendrocytes (OLs) is a notoriously difficult technique and often displays variable efficiency and cellular heterogeneity.
- We engineered a novel reporter line carrying the fluorescent protein mOrange under the control of the OL-specific transcription factor SOX10 to track, purify and characterize OLs.
- By experimenting with diverse differentiation media, we improved the generation of SOX10-positive cells. Consequently, these cells exhibited increased consistency and effectiveness in evolving into myelinating OLs.

## Introduction

Oligodendrocytes (OLs) are a type of glial cell found in the central nervous system (CNS) that play a crucial role in the formation and maintenance of myelin sheaths, which insulate and protect axons. They arise at late stages in early brain development from specific precursors called oligodendrocyte precursor cells (OPCs). Both OLs and OPCs are essential for proper neuronal communication and the overall integrity of the nervous system ^1–3^. They have been implicated as a key drivers for disease pathology in a number of neurodegenerative disorders ^4–13^, including multiple sclerosis (MS) as characterized by the progressive loss of myelin, and a number of leukodystrophies, genetic disorders that affect the cerebral white matter. Current therapeutic approaches for white matter diseases are limited, emphasizing the urgent need for strategies that can promote remyelination and restore neurological function. Animal models and primary cultures of human OLs developed over the years have been valuable tools and models to recapitulate, at least in part, the pathological hallmarks of the diseases. Nevertheless, major limitations of these models are represented by the partial overlap of the former with humans, and the scarce availability of the latter, highlighting the need for a robust, reproducible *in vitro* human model.

Induced pluripotent stem cells (iPSCs) have revolutionized the field of cellular biology and regenerative medicine. They offer a promising avenue for developing patient-specific disease models, personalized medicine, and cell-based therapies ^14^. These pluripotent cells can differentiate into any number of cell types under the right conditions, including but not limited to neurons, astrocytes, and microglia, thus providing a renewable source of cells for disease modeling and drug discovery. In the last decade, there has been a growing interest in harnessing iPSC technology to generate glial cells, including OLs, from these cells. By recapitulating the development of human OLs *in vitro*, we can shed light on the molecular mechanisms governing the differentiation and maturation of these otherwise difficult to obtain human cells. Understanding the intricate mechanisms involved in OPC proliferation, migration, differentiation, and maturation can offer valuable knowledge for tackling hypomyelinating and demyelinating diseases, where myelin deposition or homeostasis is altered, respectively. Moreover, aside from being just a renewable source of mature OLs, the OPCs are believed to play a pivotal role in neuronal communication and plasticity, making them relevant to study neurodevelopmental disorders and brain development ^15–17^.

To date, several protocols have described how to generate OLs from iPSCs, typically involving either the overexpression of OL transcription factors to directly differentiate them from the source material ^18,19^ or a stepwise process using growth-factors and signaling molecules mimicking embryonic development ^20–23^. Directed differentiation protocols, such as reprogramming through overexpression of lineage-specific transcription factors, offer improved reproducibility and an accelerated timeline to achieve mature OLs expressing markers such as myelin basic protein (MBP). While this technology provides immense promise for studying function and myelination of terminally mature OLs (such as in the case of adult-onset disorders), it fails to fully recapitulate the developmental trajectory of the OL lineage, therefore limiting its utility in studying developmental white matter disorders and the role of the OPCs in disease pathology and therapeutics. Conversely, growth factor-based protocols provide a more physiologically relevant system, however, these methods face challenges from low efficiency to high heterogeneity amongst the generated cells, and incomplete functional maturation. During development, astrocytes and OLs arise from a common progenitor, that differentiates either directly into immature astrocytes or indirectly into myelinating OLs through the generation of OPCs. The balanced and well-timed regulation of the dual fate for these bipotent progenitors has been the object of intense discussions, particularly regarding the specific conditions affecting the correct balance between the two daughter populations ^24–26^. As a result of these overlapping developmental trajectories, one of the current limitations in obtaining OPC and OL cultures *in vitro* is the spontaneous generation of contaminant astrocytes that, while providing functional support to maturing OPCs, can influence the yield of mature OLs by rapidly proliferating and taking over the culture.

To address these limitations, we developed a novel SOX10-P2A-mOrange iPSC reporter line (SOX10^mO^), engineered to express the fluorescent protein mOrange under the control of the human SRY-box transcription factor 10 (SOX10), a critical regulator of OL specification and differentiation. By leveraging this reporter system, we can systematically track SOX10 expression dynamics during OL differentiation, providing the means to readily quantify the emergence and maturation of OPCs and of mature OLs. This real-time monitoring enhances our understanding of the underlying mechanisms governing OL development, allows purification techniques enriching for SOX10-expressing cells and provides a foundation for optimization of differentiation protocols. Moreover, the use of a SOX10-driven fluorescent reporter enables the purification of highly committed OPCs, capable of differentiating into myelinating OLs; thus, serving as an *in vitro* model for relevant studies involving the function of OLs.

With this reporter cell line, we then tested the most widely cited generation protocol for OLs ^23^ and explored different approaches and workflows to potentially ameliorate the generation of OPCs and OLs. Interestingly, through tests using different differentiation media, we identified conditions that could enhance the overall yield of SOX10-positive cells. These cells subsequently underwent differentiation into functionally mature OLs, highly committed to express MBP, under the right conditions, and demonstrating a remarkable ability to myelinate poly-L-lactide fibers in an *in vitro* setting.

Here, we present a novel workflow to generate myelinating OLs consistently and efficiently from iPSC-derived OPCs. This protocol not only extends from established methods but also offers a robust platform for producing high amounts of mature OLs essential for performing functional and high-content assays using iPSC-derived oligodendrocytes.

## RESULTS

### Generation and Validation of the SOX10-P2A-mOrange Reporter Line

To optimize the efficacy of one of the most widely used published differentiation protocols for generating OPCs and OLs ^23^, we generated a SOX10-P2A-mOrange reporter (SOX10^mO^) human induced pluripotent stem cell (iPSC) line using an in-house healthy control line (a parental line referred to as 3450) ^27^. The construct generated with CRISPR-Cas9 (Figure **1A, B**) carries the sequence of the fluorescent protein mOrange ^28^ after the coding sequence of the OL-related transcription factor *SOX10,* separated by a short sequence encoding the self-cleaving peptide P2A. The latter ensures the post-translational cleavage of mOrange results in two separated proteins: the native SOX10 which is imported into the nucleus to perform its transcription factor activity and the fluorescent protein mOrange that remains in the cytoplasm. The latter, once excited, emits in the spectrum with a maximum of 562nm. We ensured by qPCR analysis that the levels of the native *SOX10* gene were comparable in the SOX10^mO^ line and in the parental control line (Figure **1D**). Next, we validated the efficiency of the generated SOX10^mO^ cell line, in parallel with the parental control (3450) iPSCs, by inducing its differentiation into OPCs and OLs using a 95-day protocol consisting of 4 main phases, proposed by Douvaras and Fossati ^23^ (Figure **1C**). In order to establish a continuous workflow, we modified the original procedure and differentiated OPCs in bulk for at least 75 days and subsequently passaged them in appropriated cell culture vessels before inducing terminal maturation. By the end of day *in vitro* (*DIV*) 75, ∼25% of the generated cells cultured in OPC Medium, were positive for mOrange (Figure **1E, F** and Figure **2B**). Immunofluorescence (IF) analysis confirmed that >96% of the cells positive for the native SOX10 protein co-expressed mOrange, confirming that the mOrange signal is an indirect but reliable marker of *SOX10* expression (Figure **S1 A, B**). Further tests, performed by FACS and IF, revealed that ∼40% of the mOrange-positive cells expressed the platelet-derived growth factor receptor alpha (PDGFRα), a receptor primarily expressed by immature OPCs (Figure **S1 C, D**). This observations suggests that although SOX10 is known to regulate PDGFRα expression ^29^, most of the mOrange-positive cells could be in an earlier phase of the development.

**FIGURE 1.**
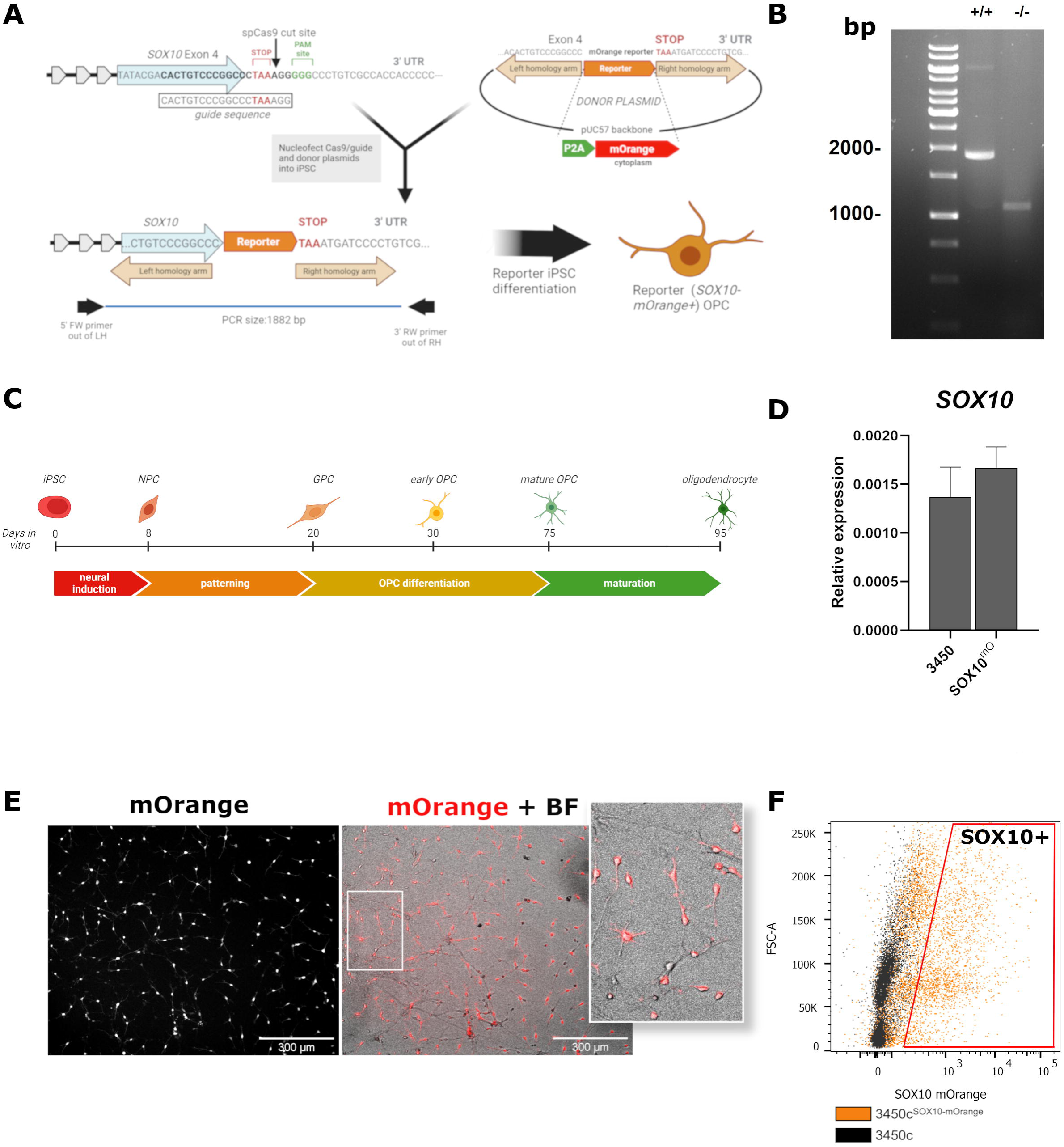
Generation of a SOX10 reporter iPSC line. (A) Schematic diagram of the mOrange reporter knock-in into the SOX10 locus using CRISPR–Cas9 genome editing. The ribonucleoprotein complex (Cas9 protein and sgRNA), and a separate “donor” plasmid containing reporter sequence flanked by homology arms were nucleofected into the parental control (3450) iPSC line. Following single-cell passaging, PCR-based genotyping was performed on individual clones using a primer set that spans the reporter sequence (both primers outside the left and right homology arms). (B) A representative PCR gel image differentiating between nucleofected (+/+) and negative control (-/-) clones. 1kb ladder is used in the first left lane for reference. (C) Representative scheme of the OL differentiation protocol used. (D) Representative histogram of a qPCR analysis at DIV75 of SOX10 gene in the reporter and parental control cell lines of the gene (n=3, one-way ANOVA). (E) Example of an OPC culture expressing mOrange driven by the endogenous SOX10 gene. Differentiation of the mOrange reporter cells was performed five times with comparable results. Scale bar: 300 μm. (F) Representative image showing the gating strategy for flow cytometry analysis of mOrange cells compared to a negative control. Black dots represent negative cells used for gating control. X axis represents mOrange.

**FIGURE 2.**
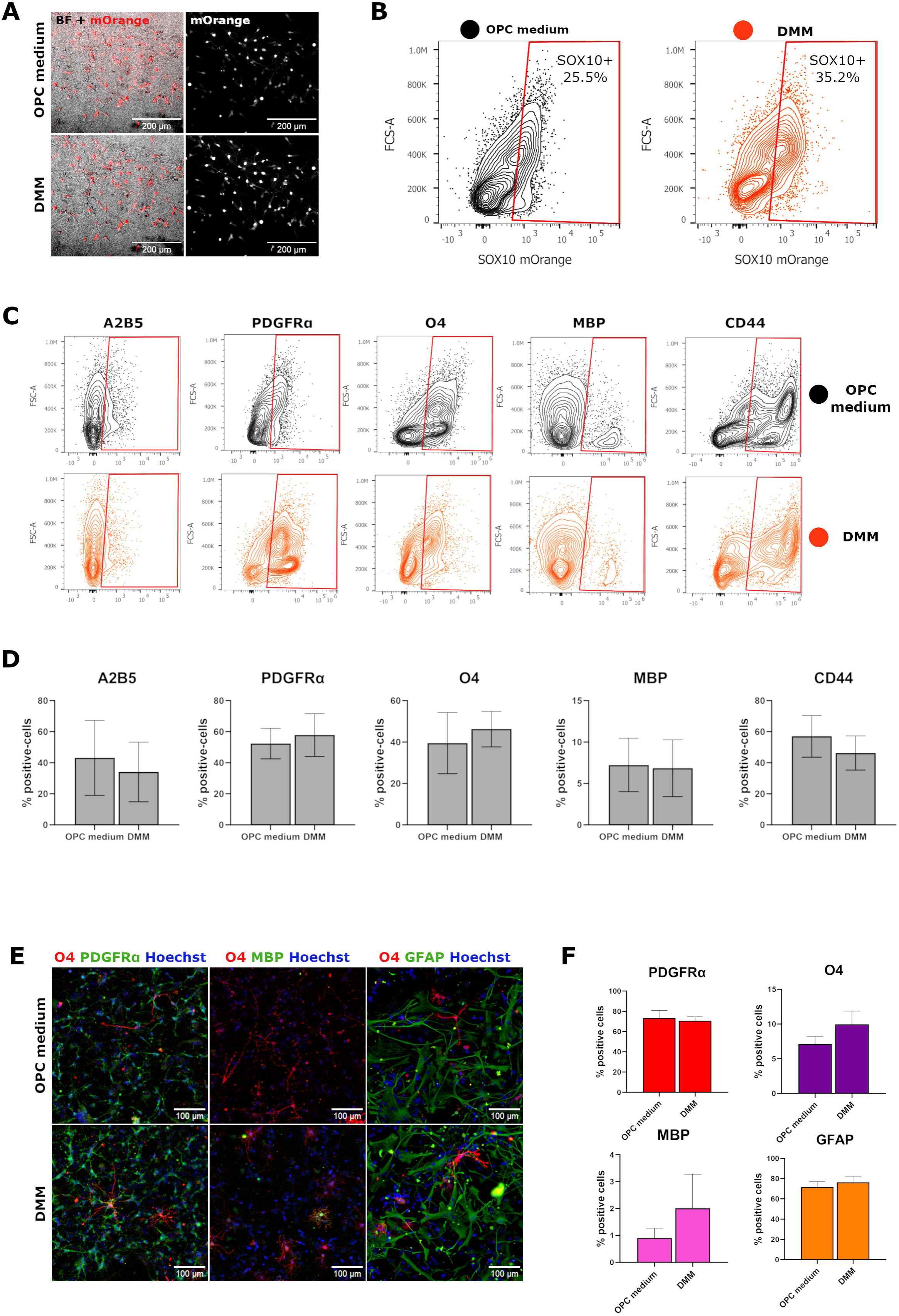
Validation of SOX10^mO^ differentiation efficiency. (A) Representative image of live cells expressing SOX10-mOrange, upper panels show cells in OPC medium, lower panels show cells in DMM. Scale bar: 300µm; (B) FACS analysis of SOX10-mOrange expression in OPC medium (left panel) and DMM (right panel); (C) FACS analysis of the lineage markers A2B5, CD44, MBP, O4 and PDGFRα. Black dots in the upper panels represent cells grown in OPC medium, orange dots in the lower panels represent cells differentiated in DMM. (D) Histograms showing the average percentages of each marker in OPC medium and DMM obtained by FACS. Error bars represent SEM (n=5, two-way ANOVA, Sidak multiple comparisons); (E) Immunofluorescence staining of cells in OPC medium (upper panel) and DMM (lower panel) showing expression of PDGFRα, O4, MBP and GFAP; (F) Histograms representing the percentage of the markers shown in panel C. Error bars represent SEM (n=3, two-way ANOVA).

### The Conventional iPSC-OL Protocol Results in Cellular Diversity and Inadequate Effectiveness in Producing Fully Functional Mature OLs

Following the Douvaras and Fossati protocol, we next switched the cells post *DIV* 75 to their maturation medium (the Glial Differentiation Medium proposed by Douvaras *et al.*, referred subsequently as ‘DMM’), to induce the maturation of the OPCs. Fluorescence-activated cell sorting (FACS) analysis of the differentiated cells following 21 *DIV* in DMM revealed a mild (∼1.5-fold) yet non-significant increase in the percentage of mOrange-positive cells (Figure **2A, B**). This finding suggests a low efficiency of DMM possibly indicative of incomplete differentiation/maturation. The observation of the mild rise in SOX10-mOrange cell percentage, upon use of DMM, and the presence of a consistent number of SOX10-negative cells led us to investigate the presence of different subpopulations of glial progenitors. To do so, we conducted a comprehensive analysis of the cell populations at each step of the protocol to gain insights into their developmental stage (Figure **2C-F**).

FACS characterization of SOX10^mO^ -positive cells highlights that, post *DIV* 75 (OPC phase) ∼50% of the cells express the early marker PDGFRα and ∼36% the ganglioside A2B5, indicative of the presence of early OPCs (see **Table 1**). When looking at glycoprotein O4, a marker indicative of a more advanced stage (late OPC/pre-OL), we found that between 20% and 40% of the total cells express this marker. This suggests that in the OPC phase of the protocol (up to *DIV* 75) a significant portion of the cells are OPCs among which a fraction is transitioning towards a more mature phase. Interestingly, a high percentage (∼60%) of the total cells expressed the surface marker CD44, a marker typical of astrocyte-restricted progenitors^30^. After addition of DMM to induce maturation of the OPCs, we observed a mild, yet non-significant, decrease of early markers such as A2B5 (from 43.6% to 38.7%) and a slight increase of PDGFRα (46.7% to 56.4%) with O4-positive cells remaining around ∼45% of the total SOX10^mO^ cells (Figure **2C, D**). Only a small fraction (∼5% of the mOrange^+^ cells) of the cells treated for 21 days with DMM expressed the mature OL marker MBP. Consistently, we still found in this phase of the protocol that 56% of the total cells (45.6% in the mOrange^+^ fraction and 10.4% in the mOrange^-^ cells) are positive for CD44, indicating a limited efficiency in generating mature OLs, with about half of the cells remaining in a late OPC phase and the other half potentially remaining committed towards an astrocytic fate.

**Table 1.** Antigenic phenotype of OL lineage cells.

| Cell Type | Positive Markers | Negative/Decreasing Markers |
| --- | --- | --- |
| Glial-Restricted Progenitors (GRPs) | A2B5, GFAP, NESTIN | PDGFRα, NG2, CD44 |
| Astrocyte-Restricted Progenitors (ARPs) | A2B5, CD44, GFAP | PDGFRα |
| Oligodendrocyte-Type II Astrocyte Progenitor Cell (O2-A) | A2B5, PDGFRα, NG2 | GFAP, CD44 |
| Oligodendrocyte Precursor Cells (OPCs) | NG2, OLIG2, A2B5, PDGFRα, CD44 | GFAP, MBP |
| Late OPCs/Pro-Oligodendrocytes (pro-OL) | NG2, OLIG2, SOX10, O4, PDGFRα, A2B5 | GFAP, MBP |
| Immature Oligodendrocytes | OLIG2, SOX10, CNP, O4, A2B5, CD44 | GFAP, PDGFRα |
| Mature Oligodendrocytes | O4, CNP, MBP, MOG | GFAP, PDGFRα, A2B5 |

### Assessment of glial progenitor markers revealed high heterogeneity

During development, OPCs and astrocytes are derived from a common progenitor, commonly known as a glial-restricted progenitor (GRP) ^2,3,24,31,32^, a tripotent cell that can give rise to type-I and type-II astrocytes as well as OPCs. This cell can undergo fate restriction to give rise to astrocyte-restricted progenitors (ARPs) expressing CD44 ^30^, or give rise to a bipotent progenitor OL/type-II astrocyte (O-2A) expressing PDGFRα ^33,34^, among other markers ^33,34^. Finally, O-2A cells can also give rise to fully committed OPCs. Moreover, all of these progenitors are known to express A2B5 ^33^, a marker that persists into the early OPC stage.

Considering those complex lineage dynamics, we next sought to interrogate our cultures to explore what percentage of GRPs is present, and to get a better understanding of the extent to which they are committed either to the astrocyte or OPC lineages. With this in mind, we identified A2B5^+^ PDGFRα^+^ cells as O-2A cells, the direct precursors of OPCs and CD44^+^PDGFRα^-^ cells as ARPs (Figure **3A**). Phenotypic analysis by flow cytometry of these populations in SOX10^mO^ cells indicates that, in OPC medium (at *DIV* 75), ∼45% of the total cells are A2B5^+^ PDGFRα^+^, consistent with an O-2A identity, while ∼18.4% of the cells expressed the CD44^+^PDGFRα^-^ antigenic profile, indicative of ARPs (Figure **3A**). Furthermore, ∼9.8% of the total cells presented with an A2B5^+^ PDGFRα^-^ profile (Figure **3A**) that we classified as tripotent GRPs. Differentiation in DMM did not significantly change the percentage of A2B5^+^ PDGFRα^+^ O-2A cells (44.5% of the total cells) with a mild but not significant increase of A2B5^+^ PDGFRα^-^ GRPs (17.3% of the total). This suggests that the addition of DMM does not change the commitment previously acquired by the progenitors. An in-depth analysis of SOX10-mOrange intensity in the aforementioned populations indicates that the A2B5^+^PDGFRα^+^ cells, identified above as O-2A cells, retain the highest intensity of SOX10, demonstrating that these are cells committed towards the OL lineage (Figure **3B**). Maturation in DMM for 21 days consistently decreased the intensity of SOX10 in this population. This observation, coupled with the previously overall mild increase of mOrange and PDGFRα percentages (Figure **2B-D**), indicates that eventually a good portion of the progenitors progress towards an OL lineage and stop expressing A2B5 and become PDGFRα^+^ OPCs. Surprisingly, ARPs, identified previously as CD44^+^PDGFRα^-^ cells, composed ∼18.5% of the total cells both in OPC medium and DMM.

**FIGURE 3.**
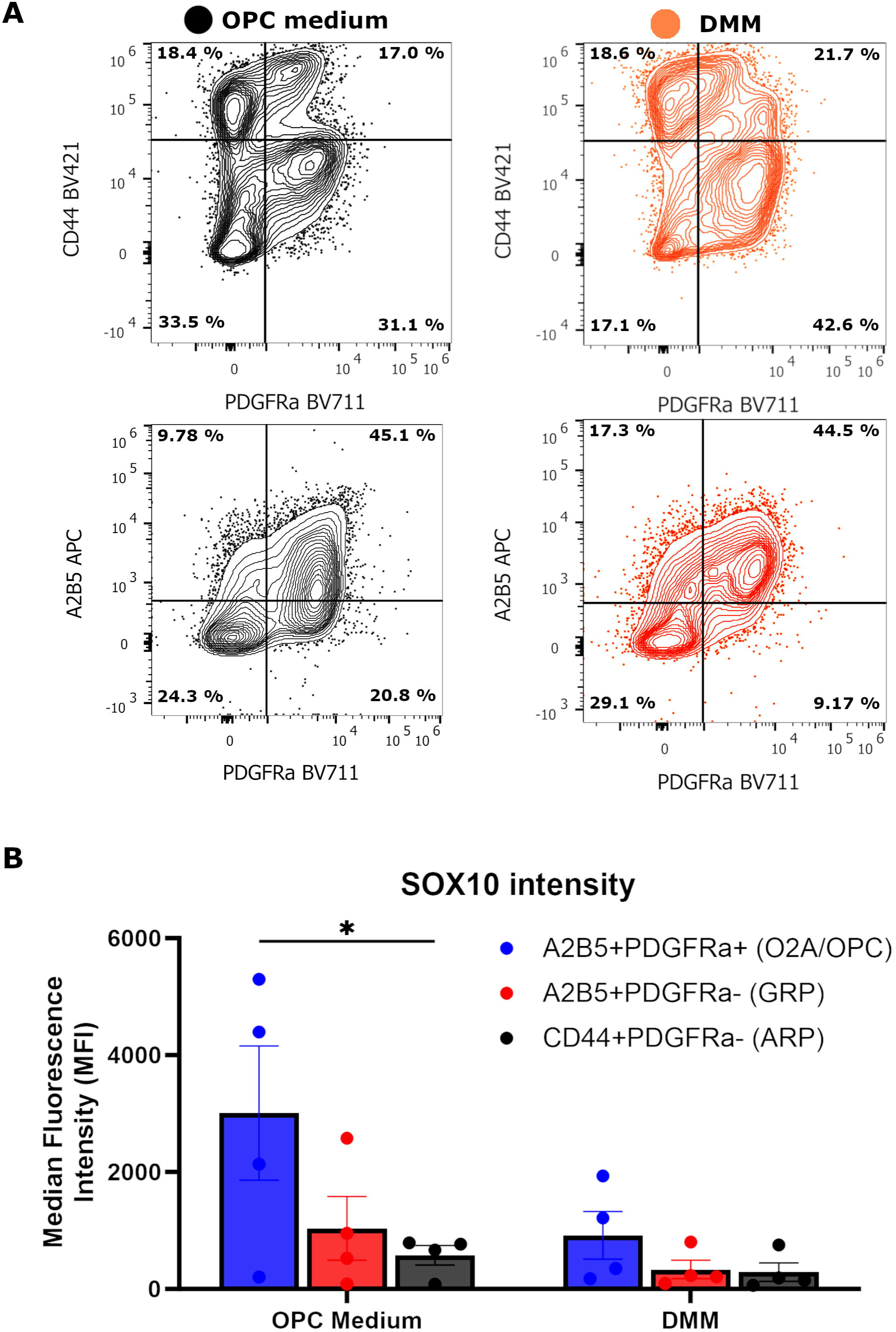
Assessment of glial-restricted progenitors. (A) Representative FACS analysis of cell populations in OPC medium and DMM. Scatterplots in the upper panels show the analysis of CD44 vs PDGFRα and lower panels show A2B5 vs PDGFRα. (B) Histogram representing the fluorescence intensity of SOX10-mOrange in each of the progenitor population identified (n=4, two-way ANOVA, Tukey’s multiple comparisons).

Altogether, this implies that, rather than a homogeneous population of OPCs, we obtained a heterogeneous cell population of GRPs with a significant fraction of them spontaneously transitioning towards an astrocytic fate. Additionally, maturation in DMM for 21 days partially enhanced the commitment of the early progenitors towards OPCs but not the terminal maturation of these cells.

### Increased Presence of Astrocytic Cells During the Maturation Phase

In agreement with our earlier findings, immunofluorescence imaging of cells after the end of the maturation phase (*DIV* 95) revealed a relatively small number of cells positive for the late OL-lineage surface marker O4 and only a sparse number of cells were detected co-expressing the myelin marker MBP, (Figure **2E, F**). In contrast, a significant proportion of these cells were glial-fibrillary acidic protein (GFAP)-positive cells both after treatment with OPC medium (*DIV* 75) and after treatment with DMM (*DIV* 95). Although GFAP is commonly regarded as an astrocytic marker, GRPs have been widely reported to express GFAP among other glial markers ^35^. We therefore concluded that either an increased number of cells undergo an astrocytic fate or the maturation phase fails to completely commit the previously obtained glial progenitors towards OL fate, as suggested by the presence of CD44- and A2B5-positive cells in the cultures.

Additional experiments conducted on the control cell lines 3450 and 81280 provided further confirmation of these findings highlighting the heterogeneous composition of the cultures. At *DIV* 95, a substantial proportion of the cell population is composed of OPCs and pro-oligodendrocytes (pro-OL, a more mature population of non-myelinating OLs), as indicated by the upregulation of OPC-related transcripts *NKX2.2*, *NKX6.2*, and *PDGFRα*. Again, examination of the total cell population at *DIV* 95, as well as throughout the differentiation protocol, shows that a *GFAP* transcript was present at high levels, further supporting the notion that a significant number of astrocytic-like cells or GRPs were present in the cultures (Figure **S5**).

Thus, we concluded that (i) the induction protocol produces a high percentage of GRPs among which there are early OPCs but also ARPs that can potentially give rise to astrocytes; (ii) that the addition of DMM for 21 days increases the percentage of committed OPCs but partially fails to bring those to terminal maturation; (iii) ARPs present already in the early stages of the protocol can proliferate and give rise to GFAP-positive astrocytes in the maturation phase. Additionally, as described in previous works ^36,37^, we successfully confirmed that a substantial number of cells retain the gene expression profile associated with a progenitor identity including but not limited to *PAX6,* a gene commonly expressed by undifferentiated neural progenitors and GRPs (i.e., bipotent progenitors).

### Novel Medium Ameliorating the Production of SOX10-Positive OPCs

Following *DIV* 75, when a majority of OPCs were anticipated, we noted the sustained high presence of cells resembling early glial progenitors. To confirm the identity of these persistent cells as early glial progenitors, we exposed them to a differentiation media known to effectively steer progenitor cells towards the astrocytic lineage according to a previously published protocol ^38,39^. The detailed procedure is described in **Materials and Methods**. We generated glial progenitors from SOX10^mO^ iPSCs with the Douvaras and Fossati protocol until *DIV* 75. Subsequently, we exposed the cultures to DMM or astrocyte differentiation media (AM+AGS). The expression of SOX10*-*mOrange was followed in cells exposed for 21 days to these two different conditions until *DIV* 95.

Surprisingly, when cultured in AM+AGS media, the cells presented with a highly significant increase in mOrange-positive cells after only 10 days of culture (Figure **S2)**, in contrast to cultures grown in the presence of DMM for the same period. After 21 days of culture (*DIV* 95), the proportion of mOrange-positive cells in both populations were analyzed by FACS. Results showed that between 40.5 and 77.8% of mOrange-positive cells were now present following incubation with AM+AGS, whereas only between 10 and 32.3 ± 12.6% of mOrange positive cells were observed in cultures grown in DMM (Figures **4C** **and** **6A**). This suggests that the glial progenitors generated in OPC medium react to the components of AM+AGS by enhancing their commitment in the OL lineage. Early works from Noble *et al.* and Martins-Macedo *et al.* show how bipotent GRPs change their fate in response to different cues, becoming astrocytes when cultured in presence of Fetal Bovine Serum (FBS)^33,34^. We sought to verify this by exposing previously obtained glial progenitors to AM+AGS supplemented with 1% FBS (FBS medium) for 21 days. An in-depth immunofluorescence analysis showed that in this condition nearly all the cells were immunoreactive for the astrocyte markers GFAP, S100β and Aquaporin-4 with little to no expression of the OL markers O4, Olig2 or PDGFRα (Figure **S3**) implying that the astrocyte-like cells were primarily ARPs that were able to proliferate and differentiate into astrocytes when FBS was present. This is consistent with previous observations showing that GRPs are committed to the astrocytic fate when cultured in the presence of FBS^33,34^ and corroborates our hypotheses on the nature of the progenitor cultures.

**FIGURE 4.**
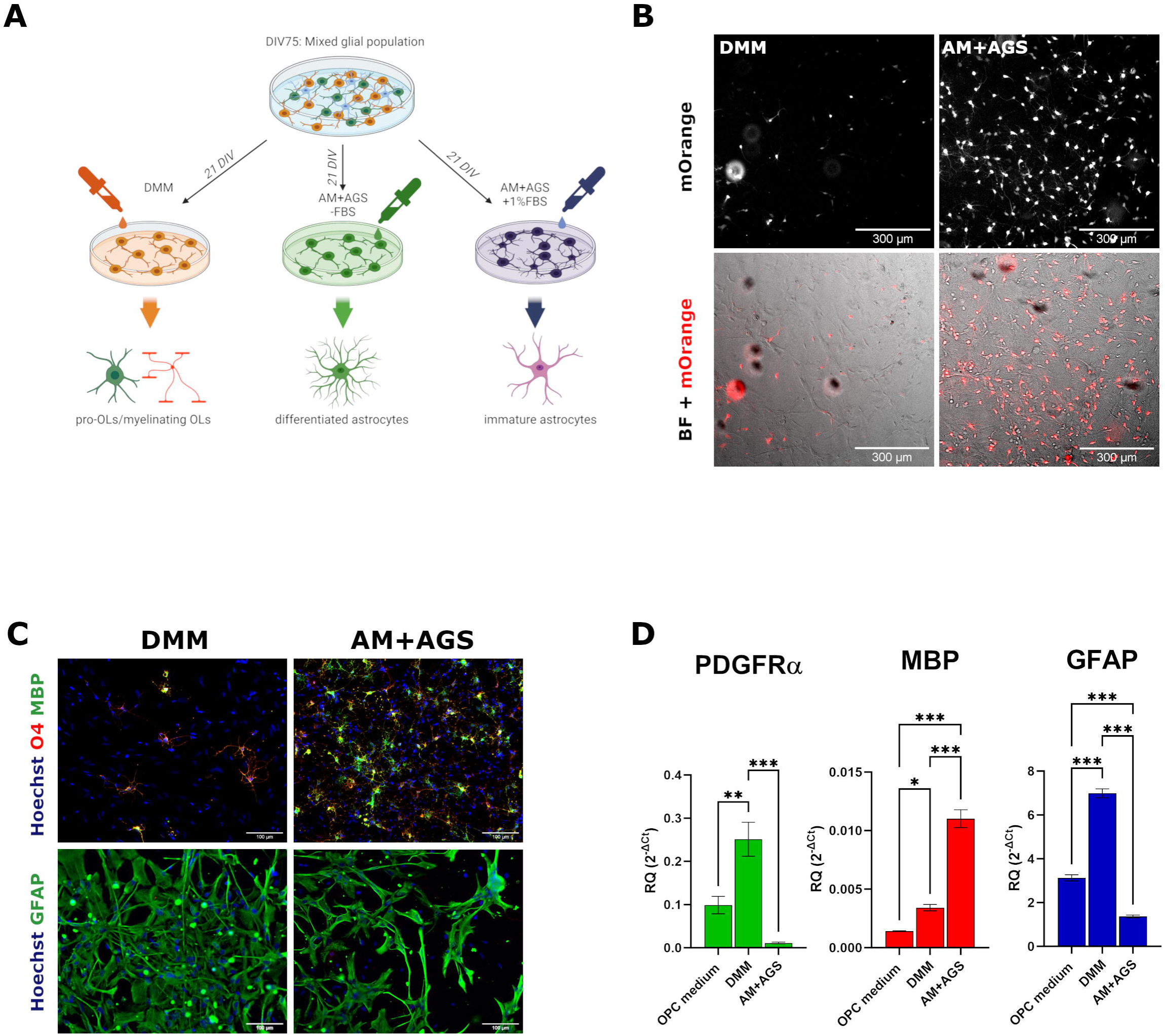
Adjustments in differentiation medium could increase the yield and maturity of SOX10+ cells. (A) Scheme representing the experimental design; (B) live imaging of mOrange expression in cells treated with DMM (left) and AM+AGS (right). Scale bar: 300µm; (C) immunofluorescence staining showing a marker comparison in cells differentiated in DMM (left panels) and AM+AGS (right panels). Upper panels show expression of oligodendroglial markers O4 (in red) and MBP (in green), lower panels show the astrocytic marker GFAP (in green). Scale bar: 100µm; (D) Histograms comparing the expression of the progenitor gene PDGFRα, the mature OL gene MBP and the astrocytic transcript GFAP in OPC medium, DMM and AM+AGS respectively. Error bars represent SEM (n=5, two-way ANOVA with Tukey’s multiple comparisons).

Taken together, we concluded that cultures exposed to OPC medium contain GRPs that under the influence of FBS and other glial differentiation cues differentiate into ARPs and then into astrocytes, but when FBS was absent, the majority of cells committed towards SOX10-positive OPCs.

### Lineage analysis of SOX10^mO^ OLs differentiated in AM+AGS

The finding that the exposure of cells to an astrocyte differentiation media lacking FBS could lead to such a drastic rise in the number of cells expressing SOX10 was unexpected but showed that the differentiation protocol could be ameliorated. This prompted us to further analyze the identity of this population and to clarify the developmental status of the cells that emerged through this workflow. Immunofluorescence staining of cells matured in AM+AGS corroborated these observations showing that the percentage of cells expressing both O4 and MBP raised by ∼2-fold compared to OPC medium, whilst displaying a highly complex and multibranched morphology, representative of mature OLs (Figures **4D** and **6C, D**). Flow cytometry analysis of cells obtained with DMM compared to cells obtained with AM+AGS revealed that the latter promoted a 1.7-fold increase in the percentage of O4-positive cells, when compared with OPC medium with numbers now reaching an average of ∼63.3% of the total population accompanied by a significant, although modest, increase in the percentage of MBP-positive cells (1.9-fold increase in AM+AGS compared with 0.7-fold in DMM, Figure **6B**). The acquisition of a more complex morphology in MBP-positive cells coincides with an increase in the intensity of mOrange (Figure **S4**), in line with the notion that SOX10 levels are positively correlated with the maturation of OLs^40^. The presence of GFAP-positive cells appeared to be sparse in cultures obtained in the presence of AM+AGS, as opposed to DMM-treated cells where their presence was far more widespread (Figure **4C**). Notably, the number of total cells in cultures exposed to AM+AGS seemed to be increased compared to DMM-treated cells suggesting the presence of mitogens in the AGS supplement that could help sustain proliferation along with other molecules that might favor OPC differentiation at the same time. Analysis of gene expression in cells differentiated in AM+AGS compared to both OPC medium and DMM, showed a robust upregulation of genes involved in the maturation of OLs such as *MBP* (Figure **4D**) and *PLP1* (Figure **S5**). Interestingly, in cells treated with AM+AGS we observed a significant downregulation in the expression of the OPC transcript *PDGFRα* as well as the astrocyte gene *GFAP* (Figure **4D**). This observation implies that, when differentiated with AM+AGS, a greater number of progenitors evolve towards an oligodendroglial fate, together with a higher number of OPCs exiting the cell cycle to complete their maturation. Similar results were obtained with two other control iPSC lines (Figure **S5 B, D**). These results show that AM+AGS is highly effective at directing greater number of progenitor cells towards an OL fate whilst at the same time helping to induce a further maturation towards myelinating OLs.

### Purification and Analysis of OPCs Expressing Different Levels of SOX10^mO^

At this point of the study, our findings substantiated a positive correlation between the level of SOX10 expression and the maturation state of the OL. This supports the notion that the expression levels of SOX10 could potentially define different subpopulations, helping us to unravel the previously described cell heterogeneity. Thus, exploiting the greater enrichment in SOX10-mOrange obtained with AM+AGS, we separated our total cell population into subpopulations based on SOX10-mOrange expression levels that could then be analyzed further as subgroups. To assess the identity of those groups, we differentiated SOX10^mO^ OPCs (*DIV* 75) with AM+AGS for 10-15 days and live-sorted the mOrange-positive cells by FACS. Based on the intensity levels of mOrange, we identified and isolated three distinct populations (Figure **5A** and **S4)**: SOX10-negative cells (SOX10^neg^), cells expressing a medium level of SOX10 (SOX10^med^) and cells expressing a high level of SOX10 (SOX10^hi^). In-bulk differentiated cells generated in parallel with AM+AGS medium were directly analyzed (non-sorted cells) along with a non-maturated control cells consisting of cells kept in the OPC medium and analyzed as a non-sorted bulk. All cells were then either collected for RNA extraction and qPCR analysis or replated onto nanofiber plates to test their ability to ensheath poly-L-lactide nanofibers. Interestingly, with cells sorted and plated onto nanofiber plates, a notable increase in mOrange-positive ensheathing cells was observed in the SOX10^hi^ sorted populations cultured in AM+AGS medium compared to the non-sorted cells grown in OPC medium, which remained in an immature state (Figure **5C**). Immunofluorescence analysis of these cells confirmed the presence of highly complex MBP-positive cells exclusively in the SOX10^hi^ sorted cells while almost no expression was detected in the SOX10^neg^ and SOX10^med^ populations (Figure **5B**). Gene expression analysis confirmed the higher degree of maturation of non-sorted cells differentiated in AM+AGS compared to non-sorted cells in OPC medium (Figure **5D**). Moreover, gene analysis of the three isolated populations highlights the superior maturity of SOX10^hi^ cells showing a strong enrichment in the levels of the maturation genes *CNP*, *PLP1* and *MBP* compared to a mild enrichment in SOX10^med^ and virtually no expression in SOX10^neg^. Flow cytometry analysis (Figure **S4**) showed that the SOX10^hi^ cell population showed a 102.6-fold increase in the intensity levels of MBP. This confirmed that OL differentiation and maturation in our cultures is sustained by a strong expression of SOX10 in a dose-dependent fashion. Consistently, the levels of the astrocytic gene *GFAP* were strongly reduced in SOX10^hi^ population compared to the non-matured control population but conversely seemed to remain enriched in the SOX10^neg^ population, in line with the notion that SOX10 expression is associated with OL commitment. Similarly, both the non-matured control cells and the SOX10^neg^ population show a similar enrichment in the early progenitor gene *PAX6.* Its levels were already decreased by 3-fold (although not statistically significant) in non-sorted cells cultured in AM+AGS compared to cells in OPC medium and is found to be strongly enriched in SOX10^neg^ cells. These findings suggest that during OPC induction phase, a significant portion of SOX10-negative cells are not yet committed to the OL fate and are likely to spontaneously generate GFAP-positive astrocyte-like cells. Analysis of the expression levels for the gene *PDGFRA*, a marker of the early phases of OPC specification, shows a greater enrichment in the SOX10^neg^ population in line with the previous observation that, at least in our cultures, SOX10 is associated with later stages of OL development rather than defining the whole OPC population. In agreement with this observation, the levels of *NKX2.2* and *NKX6.2*, two genes associated with later stages of OL maturation^41,42^, are proportionally increased in SOX10^med^ and SOX10^hi^ populations compared to SOX10^neg^. A similar observation was made for the pan-OL gene *OLIG2,* whose levels are significantly increased in SOX10^med^ and SOX10^hi^ populations but decreased in SOX10^neg^ cells. Intriguingly, when we analyzed the levels of the transcript *mKI67,* encoding the proliferation marker Ki-67, we found its levels significantly elevated in the SOX10^med^ and SOX10^hi^ populations indicating that cells expressing high levels of SOX10 are still highly proliferative (Figure **5D**). Thus, despite unequivocally exhibiting markers indicative of advanced differentiation and maturation, the SOX10^hi^ cells obtained in abundance through the utilization of AM+AGS medium retain a remarkably high degree of proliferation. This observation is intriguing, as these cells might be expected to display a reduction in their proliferative propensity at this stage. Overall, cell sorting provided the means to enrich for the SOX10^hi^ population compared with both SOX10^neg^ and SOX10^med^, helping to confirm a strong correlation between SOX10 expression and the ability to attain a terminal stage of maturation, as already established previously^29,40^.

**FIGURE 5.**
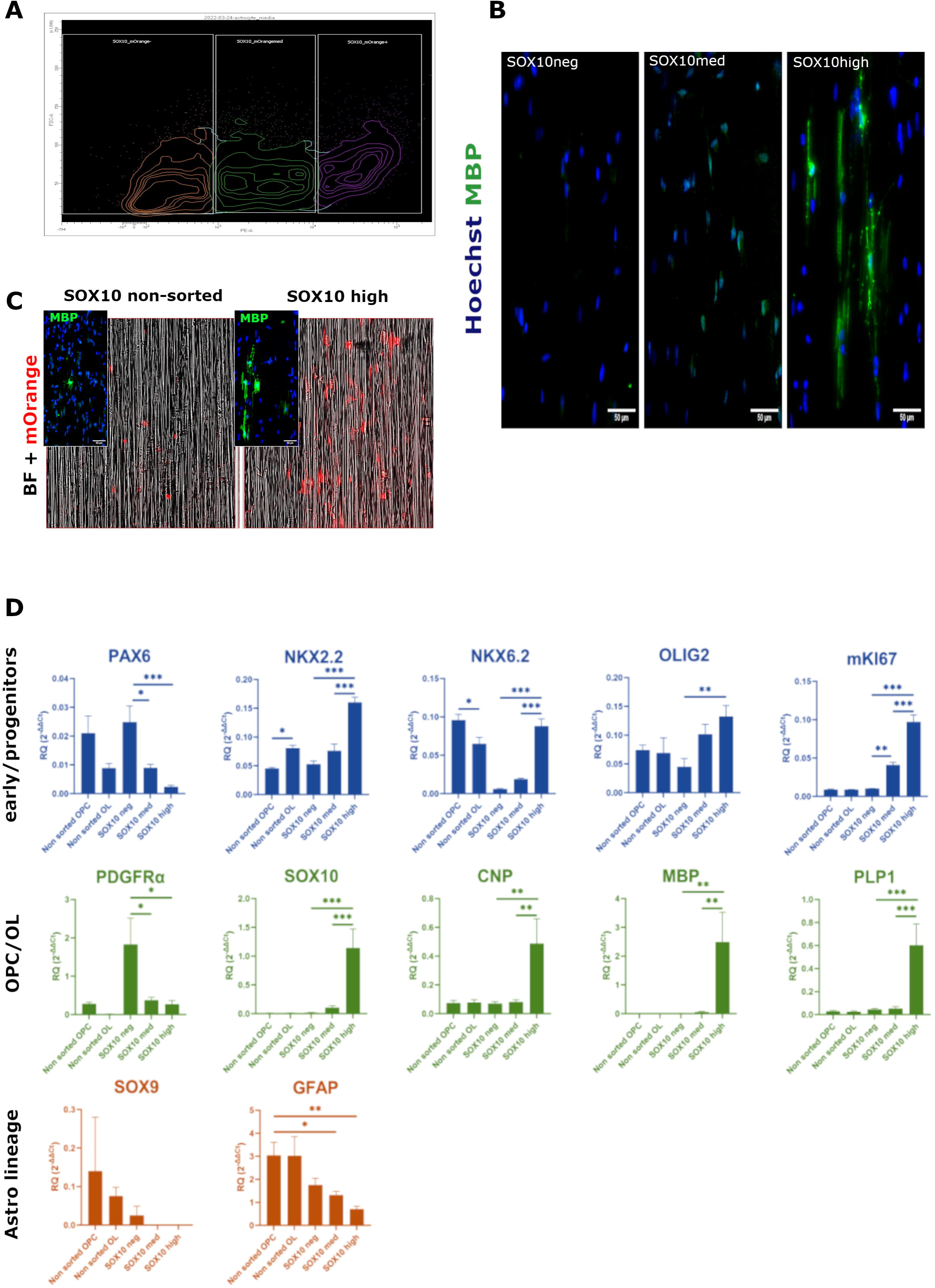
Analysis of purified SOX10^mO^ cell populations. (A) The three populations are separated by FACS based on the intensity of mOrange expression; (B) Immunofluorescence showing the isolated populations plated onto poly-L-lactide nanofibers. Myelination is assessed through staining with anti-MBP antibodies (in green), nuclei are detected with Hoechst 33342 (in blue). Scale bar: 50µm; (C) Comparison of mOrange expression in non-sorted cells vs SOX10 high sorted cells. MBP staining (in green) is shown in the insert boxes; (D) Real-time quantitative PCR analysis of the purified cells. Asterisks represent significance (n=4, one-way ANOVA with Tukey’s multiple comparisons).

**FIGURE 6.**
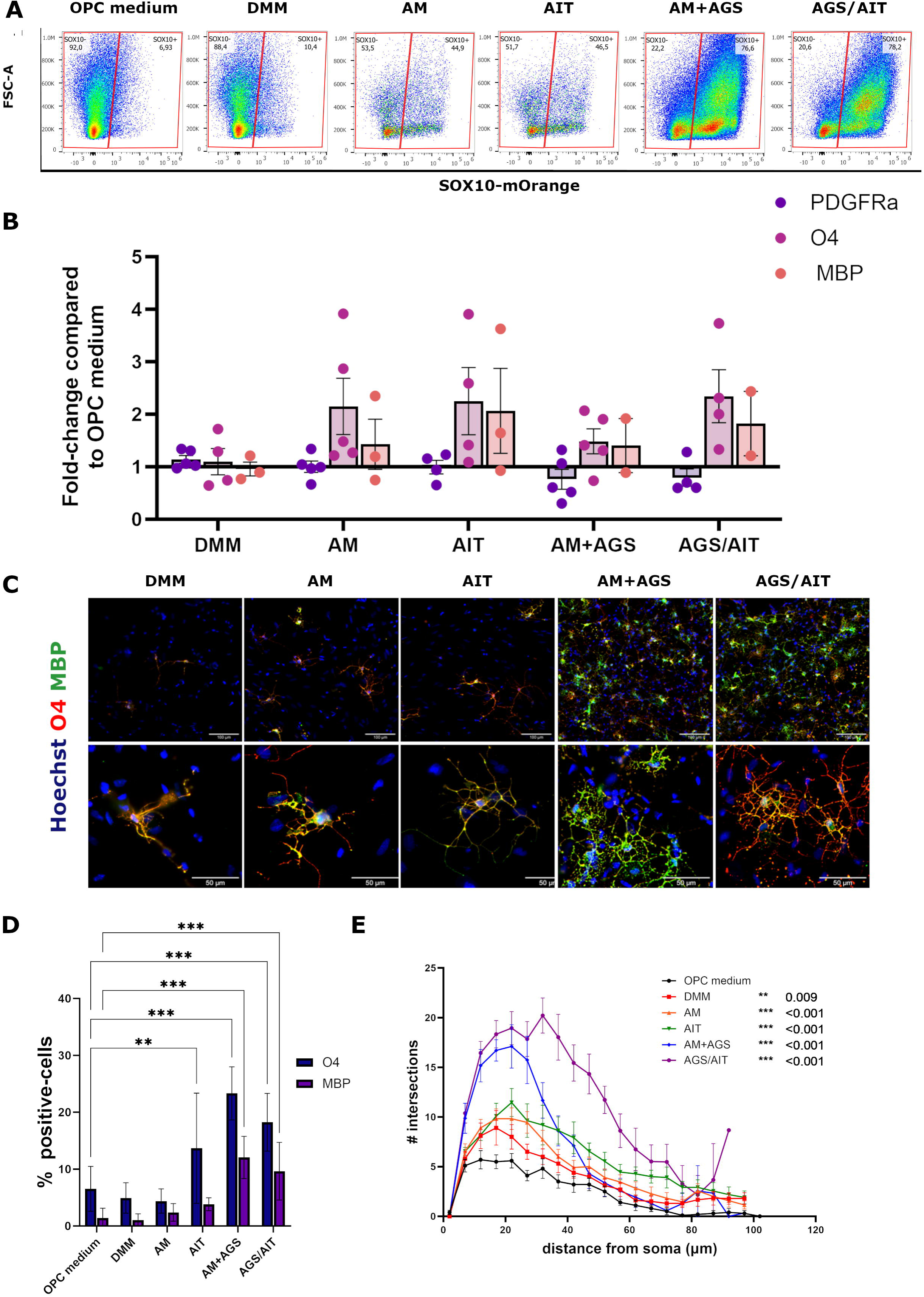
Improving maturation conditions in SOX10^mO^ cells. (A) FACS quantification of mOrange-positive cells in different culture conditions; (B) Histogram representing fold-change of flow cytometry quantifications for OL lineage markers in each maturation medium compared with OPC medium (n=5); (C) Day 95 OPCs in each maturation medium staining O4 (in red) and MBP (in green) at 20X (upper panels, scale bars:100 µm) and 63X (lower panels, scale bars: 50µm); (D) IF quantification of percentage of O4+ (blue bars) and MBP+ cells (purple bars) at day 95 in each maturation medium. Error bars represent SEM, asterisks represent significance (n=5, two-way ANOVA with Tukey’s multiple comparisons); (E) Sholl analysis quantification depicted as mean number of intersections at each ring. Error bars represent SEM, significance and p-values are shown in the graph legend (n=10, two-way ANOVA with Tukey’s multiple comparisons).

### Improvement of the OL Maturation Phase

The high proportion of proliferating SOX10^hi^ cells revealed by the previous experiments prompted us to hypothesize that the molecular cues contained in the supplement AGS stimulate the rapid proliferation of immature progenitors and might hinder their complete maturation. This could result in cells being trapped in an intermediate state between early OPCs and pro-OLs. Thus, we sought to test whether the withdrawal of the supplements could force a higher number of cells to exit the cell cycle and to complete the maturation. With this in mind, we assessed the effect of various combinations of AM basal medium, with and without supplementary components.

SOX10^mO^ OPCs generated at *DIV* 75 were cultured for 21 days across these different media conditions and compared to DMM. Assessment of the expression of mOrange through FACS revealed that AM alone was sufficient to raise the percentage of mOrange+ cells to 50.8 ± 6.3% compared to 20.7 ± 9.5% in DMM (Figure **6A**). In addition, we tested AM supplemented with T3 and IGF-1 (AIT medium), two factors proven to stimulate the maturation of OLs^3,43–47^. This condition slightly increased the percentage of mOrange-positive cells (57.9 ± 14.9%) but the highest percentage was obtained with the addition of AGS (77.8 ± 4.9%), as reported above. From these findings, we concluded that the AM media alone serves as a robust inducer of SOX10-positive OPCs, and the inclusion of AGS not only amplifies the proliferation of SOX10-positive OPCs but prompts a greater proportion of these cells to spontaneously exit the cell cycle in comparison to other conditions, a phenomenon that we potentially attributed to contact inhibition.

Following all the insights gained through these tests, we explored how integrating these findings could help in optimizing the differentiation efficiency of OLs through a two-step strategy.

First, starting from OPCs at *DIV* 75, we induced the maturation and proliferation of SOX10-positive precursors with AM+AGS supplemented for 10 days. We previously observed that by this time, a peak in the expression of SOX10 is observable along with a consistent proliferation rate (Figure **S3**). Second, we withdrew AGS and supplemented the AM basal medium with T3 and IGF-1 (AGS/AIT) until day 21 (*DIV* 95) to further induce the maturation of the SOX10-positive pre-oligodendrocytes. This experimental strategy was performed both on SOX10^mO^ cells and in a number of other control cell lines. We chose the cell line 3450, the parent cell of SOX10^mO^ regularly used in our laboratory as an in-house control^27^ cell lines and the cell line 81280 (Figure **S5**). As reported in Figure **6A** and **6B**, FACS lineage analysis revealed that this method yields a comparable percentage of SOX10-mOrange+ cells (78.2%); a higher percentage of MBP+ cells (12.5%) compared to 7.8% obtained in AM+AGS, confirming a higher degree of maturation. In addition, we calculated the percentage of O4^-^MBP^+^ cells, to evaluate the degree of maturity of the cells and we found an elevated percentage of these in AGS/AIT compared to other culture conditions examined. Similar results were obtained in control cells with FACS, IF and qPCR analyses (Figure **S2**). Immunofluorescence analysis (Figure **6C, D**) confirms FACS observations with a visible increase in the percentage of O4 and MBP-positive cells in both AM+AGS and AGS/AIT with a significantly higher percentage of MBP in this last condition. Finally, to better characterize the level of maturity of the OLs generated, we considered the morphological evaluation of cell branching, a well-recognized indicator of maturity of OLs in the field. We used Sholl analysis^48^ to calculate the degree of complexity of OLs in the different conditions. We found a higher degree of complexity, expressed in number of internodes at 10 and 20 µm from the cell body in AGS-AIT compared with AM alone suggesting a superior maturity achieved by the exposure to the two well-characterized growth factors (Figure **6E** **and S6**).

Altogether, these data showed a substantial improvement in the commitment of GRPs and in the maturation of iPSC-derived OLs.

## Discussion

In the present work, we address the delicate problem of the differentiation and maintenance of human OL lineage cells from iPSC. We focused on reducing cell heterogeneity and increasing efficiency the currently known physiological differentiation methods. Using state of the art biomolecular techniques (such as CRISPR-Cas9 genetic modification), we obtained a SOX10-P2A-mOrange reporter line that allows us to track, purify and study committed populations of iPSC-derived OPCs obtained *in vitro*. Using this novel tool, we were able to test one some of the most widely used differentiation methods and to analyze, through precise lineage-specific markers, the cell populations obtained. We demonstrate, both in the OPC phase of the protocol and at later stages, the presence of a non-negligible percentage of glial-restricted progenitor cells (GRPs), the common precursors of astrocytes and OLs. As previously described by Noble *et al*.^33^, these cells have the potential (*in vitro*) to differentiate into astrocytes in the presence of FBS or into OLs with a defined serum-free medium. Additionally, they can undergo spontaneous differentiation under conditions that are not well understood. We concluded that (i) this could be the origin for contaminant astrocytes that are often observed in these cultures and that (ii) we could exploit this nature of the progenitors to push them towards becoming astrocytes or OLs as needed. Regarding our first point (i), further studies will be needed to understand how to restrict the fate of GRPs to OLs exclusively. As for (ii), we used a method published by our lab, using a commercial medium in the presence or absence of FBS to restrict the fate of these precursors to astrocytes or OL respectively. As expected, the presence of FBS resulted in a transient upregulation in the number of SOX10-mOrange positive cells followed by a consistent increase in the percentage of GFAP-positive astrocytes. Surprisingly, in absence of FBS, the combination of astrocyte medium (AM) with its commercial supplements (AGS) led to a major increase in the percentage of SOX10-positive cells, suggesting that with this condition, more cells move towards an OL fate. Subsequent analyses indicated that the cells obtained with this medium are indeed mature OLs, expressing among the other markers, the surface protein O4 and the myelin basic protein MBP, typical of myelinating OLs in a significantly higher percentage compared to the previous protocols. Although OPCs are typically described as cells with a certain degree of self-renewal, those obtained from iPSCs with existing methods partially lose their ability to respond to the differentiation media when subjected to passage or frozen and thawed (data not shown). However, even after 5 passages or a freeze-thaw cycle, the AM+AGS medium showed a consistent efficiency in producing a high percentage of SOX10-positive OPCs. This is a remarkable finding as it opens the possibility of maintaining a large number of iPSC-derived OPCs in culture for longer periods of time or to be cryogenically stored without having to differentiate until needed. Another interesting observation is the increased proliferation rate displayed by glial progenitors cultured in AM+AGS. Developing a rationale to explain this observation has been challenging due to the unidentified composition of the AGS supplement. The simultaneous occurrence of this phenomenon alongside a significant rise in the percentage of SOX10-positive cells prompted us to inquire whether this could be the result of different cellular subpopulations responding differentially to unknown components within the medium. FACS analyses of OPCs subjected to AM+AGS show a scattered distribution of SOX10-positive cells. Based on the intensity of mOrange, three populations could be identified and transcriptionally characterized: (i) a SOX10-negative population enriched in genes typical of early glial progenitors such as *PAX6* and *PDGFRA*, but also *GFAP,* a gene found to be upregulated not only in astrocytes but in GRPs. We interpret this as the most undifferentiated and heterogeneous cell population. (ii) An OPC population, that starts to express SOX10 at mild levels, displaying a significant increase in lineage specific genes such as *OLIG2* along with maturation-associated transcripts such as *NKX2.2* and *NKX6.2*. This is identified as a more homogeneous population of late OPCs that is starting to upregulate maturity genes such as *PLP1* and *MBP*. Finally, (iii) a population expressing high levels of SOX10, highly enriched in maturation genes such as *CNP*, *PLP1* and *MBP.* This is the only population that once plated onto nanofibers displayed the ability to produce MBP-positive myelin sheaths and therefore is enriched in functionally mature OLs.

Remarkably, this is also the population that showed the highest upregulation of *mKI67*, the gene encoding for the proliferation marker Ki-67. This observation is puzzling as no proliferation would be expected in such a differentiated cell pool and suggests that there may be one or more mitogens in the media and supplement formula that prevents cells from exiting the cell cycle and reaching terminal maturation. Further cell cycle studies will be needed to test this idea. With this in mind, we improved the differentiation of the cells by controlling the composition of the medium: the absence of AGS supplements prevented rapid proliferation (not shown) but did not lead to a significant upregulation in differentiation markers. Moreover, substitution of AGS for the well-known OL-associated trophic factors IGF-1 and T3 significantly improved the percentage of MBP-positive cells. From this observations and findings, it ultimately led us to a two-step maturation strategy: first, we induced proliferation of SOX10-positive OPCs by exposing them to AGS. Once the percentage of SOX10-positive cells increased (after 10 days of treatment), we withdrew AGS and induced maturation of the SOX10 cells with the addition of IGF1 and T3 for additional 10 days. This resulted in a mild, yet significant increase in the percentage of MBP-positive cells, displaying a complex morphology typical of mature OLs, as corroborated by Sholl analysis of these cultures. However, it remains to be clarified why not all the SOX10-positive cells gave rise to myelinating OLs. This may be a consequence of the culture duration or more stringent conditions may be necessary?

In the present work, through the use of a SOX10-P2A-mOrange reporter iPSC line, we analyze the complex lineage relationships that are at the origin of glial progenitors’ heterogeneity. This information was used to improve the commitment of glial progenitors towards the OL fate and to develop a two-step maturation approach. This helped us to achieve a more consistent and highly efficient workflow to increase the yield of mature OLs.

In conclusion, using a novel SOX10-P2A-mOrange reporter line, we improved an existing differentiation method to provide a highly reproducible workflow to differentiate iPSC-derived glial progenitors into mature OLs, whilst explaining why heterogeneity would often arise with these glial cultures. This protocol enabled the generation of cultures highly enriched in SOX10-positive OPCs that can be matured into MBP-positive OLs. However, further efforts will be necessary to refine the differentiation process, ensuring a complete commitment to the oligodendrocyte lineage that effectively recapitulates every phase of their development. Taken together, these findings lay the foundation for a robust two-step workflow that can be used to develop highly pure, and mature cultures of OPCs and OLs for discovery and translational studies.

## List of abbreviations

AM: astrocyte medium
ARP: astrocyte-restricted progenitor
BSA: bovine serum albumin
CNS: central nervous system
FBS: fetal bovine serum
GFAP: glial-fibrillary acidic protein
GRP: glial-restricted progenitor
IF: immunofluorescence
iPSC: induced pluripotent stem cell
MBP: myelin basic protein
MS: multiple sclerosis
O2-A: oligodendrocyte-type II astrocyte progenitor cell
OL: oligodendrocyte
OPC: oligodendrocyte precursor cell
PFA: paraformaldehyde
SOX10: SRY-box transcription factor 10.

## Materials and Methods

### Cell lines

The lines used were: 3450^27^ and 81280 (Genome Quebec). The complete profiles of the iPSCs are listed in Table 2. The use of iPSCs and stem cells in this research was approved by the McGill University Health Centre Research Ethics Board (DURCAN_IPSC/2019-5374).

**Table 2.** Overview of hiPSC.

| Cell line | Cell Type | Donor's Age | Sex<sup>1</sup> | Cell Source | Reprogramming | Media<sup>2</sup> |
| --- | --- | --- | --- | --- | --- | --- |
| 3450 | PBMC | 37 | M | C-BIG | Episomal | E8 |
| 81280 | Fibroblasts | 50 | M | Genome Quebec | Sendai | mTeSR1 |

<sup>1</sup> M, male; F, female. <sup>2</sup> Optimal media used for cell growth.

### Generation of SOX10*-*P2A-mOrange reporter line

The SOX10-P2A-mOrange reporter cell line was generated using the iPSC line 3450 through CRISPR-Cas9 method as previously described ^49^.

Briefly, the sgRNAs were designed via the CRISPR Design Tool https://www.benchling.com with the following sequence: CACTGTCCCGGCCCTAAAGG and synthesized by Synthego. For the SOX10 donor template, left homology arms + P2A-mOrange + right homology arm cassette was synthesized by Bio basic company, then cloned into pUC51 plasmid. Alt-R® S.p. HiFi Cas9 Nuclease V3 was purchased from *IDT*. For gene editing at the SOX10 locus, gRNA/Cas9 RNP complex along with SOX10 donor template plasmid were nucleofected into 3450 iPSC (*Lonza Biosciences*). After nucleofection, cells were seeded into 96 well plate. Edited cells were screened by PCR-based genotyping with 5’OUTF and 3’OUTR primers. PCR analysis and sanger sequencing confirmed successful P2A-mOrange cassette integration in both chromosomes.

### Oligodendrocyte and astrocyte differentiation

Oligodendrocyte differentiation was adapted from a previously described method ^23^. Briefly, the protocol consists of three main phases: in the first phase, iPSCs are induced towards neural progenitors for 8 DIV using SB31542 (10µM) and DMH-1 (2µM, *Selleck Chemicals LLC*), patterned with Retinoic Acid (100nM, *Sigma-Aldrich*) and Purmorphamine (1µg/mL, *Sigma-Aldrich*) for 4 days in monolayer and then grown in suspension in the same medium for 8 days to form spheres. In the second phase, OPCs are generated and expanded in a suspension culture with the addition of OPC medium, supplemented with PDGF-AA (10ng/mL, *Peprotech*), IGF-1 (10ng/mL, *Peprotech*), NT-3 (10ng/mL, *Peprotech*), HGF (5ng/mL, *Peprotech*), 3,3,5-Triiodo-l-thyronine (T3, 60ng/mL, *Sigma-Aldrich*), Biotin (100ng/mL, *Sigma-Aldrich*), and Dibutyryl cAMP (db-cAMP, 1µM, *Carbosynth*) for 10 days. Subsequently, the cells are grown onto poly-L-ornithine/laminin-coated vessels (*Sigma-Aldrich, Invitrogen*) to complete the differentiation towards an OPC lineage until *DIV* 75. At this point cells can be detached and passaged using TrypLE™ Express Enzyme (*Thermo-Fisher*) or frozen. For the third and final phase, 5000 cells/cm^2^ are plated onto poly-L-ornithine/laminin-coated vessels and cells are allowed to complete their differentiation in a medium (DMM) supplemented with Ascorbic Acid (20µg/mL, *Sigma-Aldrich*), Biotin, T3 and db-cAMP.

Astrocyte differentiation was performed by modifying a previously described protocol ^38,39^. Briefly, progenitors differentiated at *DIV* 75 of the Fossati protocol, undergo an initial phase of induction and expansion towards an astrocytic lineage with the addition of Astrocyte Growth Medium (referred here as ‘AM’) supplemented with astrocyte growth supplements (named here as ‘AGS’, *ScienCell Research Laboratories*) or AM and AGS supplemented with 1% of Fetal Bovine Serum (Thermo-Fisher) for 21 days. Modified differentiation media were obtained by using AM alone, AM supplemented with IGF-1 (*Peprotech*) and T3 (*Sigma-Aldrich*) or AM supplemented with AGS for 21 days. In one condition cell cultures passaged after *DIV* 75 were differentiated in AM+AGS for 10 days and allowed to mature in AM supplemented with IGF-1 and T3 (AIT medium) for 11 days.

### FACS analyses and live sorting

Live sorting was performed using the Aria Fusion cell sorter (*BD Biosciences*). To collect cells for analysis by flow cytometry, TrypLE™ Express Enzyme (*Thermo-Fisher*) was added to each well and incubated at 37℃ for 10 min. Samples were quantified on the Attune flow cytometer (*Thermo-Fisher*), to determine the volumes to be used for staining. LIVE/DEAD™ Fixable Aqua Dead Cell Stain kit (*Thermo-Fisher*) was used to stain for viability, according to manufacturer instructions. Using the Human TruStain FcX™ antibody (*BioLegend*), samples were blocked according to manufacturer instructions. To prepare the pre-conjugated antibodies, each was diluted in FACS buffer (D-PBS, *Wisent* & Fetal Bovine Serum, *Thermo-Fisher*), and appropriate controls were prepared as well. Antibodies used for staining are indicated in the table below.

### Immunofluorescence

To analyze cells by immunofluorescence staining and microscopy, the 96-well clear bottom black cell culture plate (Falcon) was used for culturing. Anti-O4 (Mouse, *R&D System*) was stained live, and incubated at 37℃ for 30 min. The cells were fixed with 2% paraformaldehyde (PFA) and permeabilized with PBS containing 0.2% Triton-X-100 (*Sigma-Aldrich*). Cells were blocked for one hour at room temperature with PBS containing 0.05% Triton-X and 5% bovine serum albumin (BSA, *Sigma-Aldrich*). The primary antibody was added according to the manufacturer-suggested concentrations diluted in blocking solution and left at 4℃ overnight (primary antibodies are listed in **Table 3**). The next day, secondary antibodies (Alexa Fluor, *Invitrogen*) were added at final concentrations of 1:400 diluted in blocking solution and incubated for two hours. Nuclei were stained for 10 minutes with Hoechst-33342 (*Thermo-Fisher*) diluted in PBS for a final concentration of 1:4000. The plate is stored at 4℃ until imaging. Plates were imaged with a 10X objective on a Thermo-Fisher CX5, or ImageXpress (*Molecular Devices*) high-content imaging platform. Detailed 63X images were obtained using a *Zeiss* AxioObserver Z1 Inverted Microscope. For images acquired with either platform, analysis was performed using one software at least, and images manually acquired were analyzed using the ImageJ software.

**Table 3.** List of antibodies used for flow cytometry.

| Antibody | Fluorochrome | Ab clone | Manufacturer (CAT#) | Dilution used |
| --- | --- | --- | --- | --- |
| A2B5 | APC | 105HB29 | Miltenyi Biotec (130-124-010) | 1:400 |
| PDGFRa | Brilliant Violet 711 | 16A1 | BD Optibuild (752901) | 1:50 |
| O4 | VioBright B515 | REA576 | Miltenyi Biotec (130-120-016) | 1:100 |
| MBP | purified* | MBP101 | Abcam (ab62631) | 1:3200 |
| CD44 | Brilliant Violet 421 | BJ18 | Biolegend (338810) | 1:1000 |

\* Antibody coupled with a PE-Cy7 fluorophore with a commercial conjugation kit (Abcam (ab102903)). Manufacturer's protocol followed.

**Table 4.**
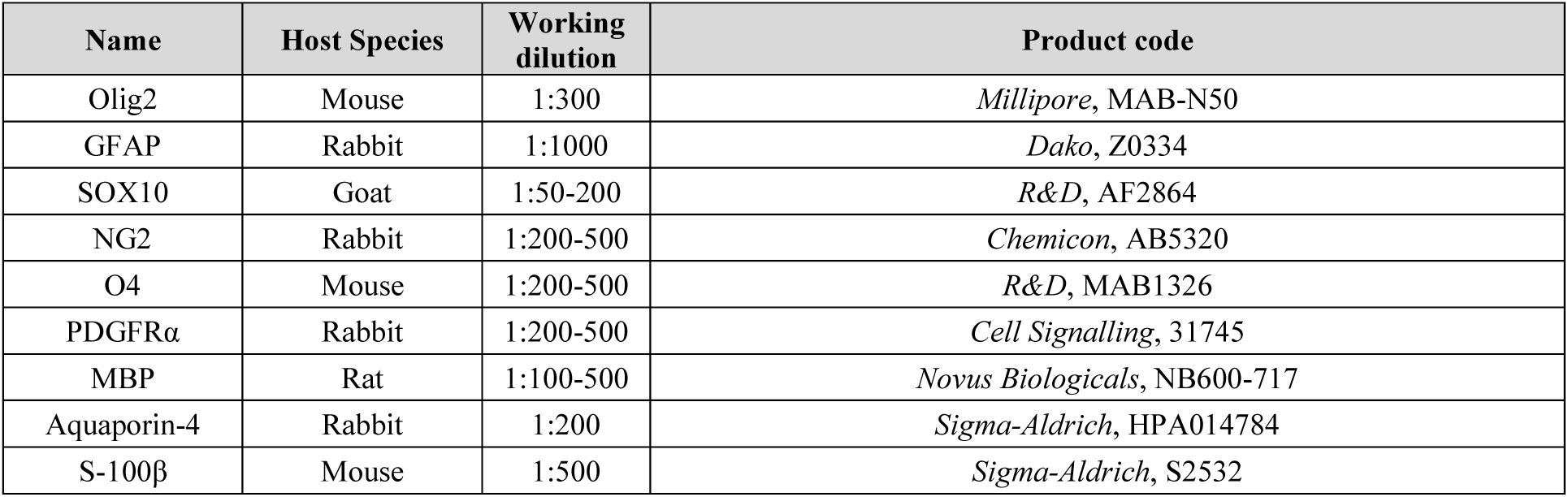
List of primary antibodies for immunofluorescence.

| Name | Host Species | Working dilution | Product code |
| --- | --- | --- | --- |
| Olig2 | Mouse | 1:300 | *Millipore*, MAB-N50 |
| GFAP | Rabbit | 1:1000 | *Dako*, Z0334 |
| SOX10 | Goat | 1:50-200 | *R&D*, AF2864 |
| NG2 | Rabbit | 1:200-500 | *Chemicon*, AB5320 |
| O4 | Mouse | 1:200-500 | *R&D*, MAB1326 |
| PDGFRα | Rabbit | 1:200-500 | *Cell Signalling*, 31745 |
| MBP | Rat | 1:100-500 | *Novus Biologicals*, NB600-717 |
| Aquaporin-4 | Rabbit | 1:200 | *Sigma-Aldrich*, HPA014784 |
| S-100β | Mouse | 1:500 | *Sigma-Aldrich*, S2532 |

### Real-time quantitative PCR

RNA was extracted according to Qiagen RNeasy RNA extraction kit manufacturer instructions. RNA was reverse transcribed using the iScript Reverse Transcription Supermix (*Bio-Rad*), and cDNA was stored at −20 until use. For expression analysis, probes against genes of interest were sourced from the *Thermo-Fisher* database (TaqMan, *Thermo-Fisher*). A complete list of probes is displayed in **Table 5**.

**Table 5.**
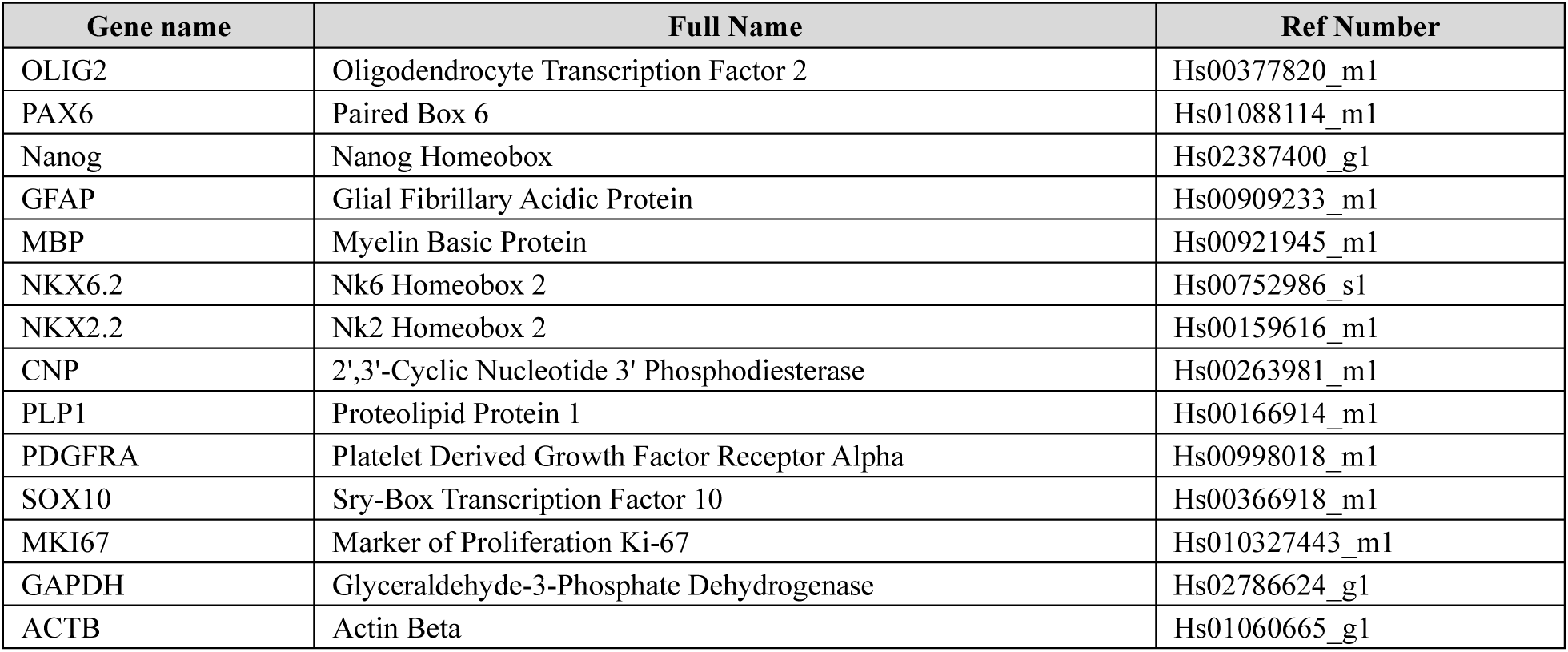
List of TaqMan probes for RT-PCR.

### Sholl analysis

For Sholl analysis, O4 stained cells (n=10/condition) were acquired a *Zeiss* AxioObserver Z1 Inverted Microscope at a magnification of 63X. Images were processed using ImageJ software as follows: single cell images were cut out and mad binary using the threshold function. Sholl analysis was performed using the Neuroanatomy tool, setting the center as the centroid of the soma and setting 2µm as starting diameter and 100µm as maximum diameter with a step size of 5µm. The results were plotted average number of intersections/radius.

### Statistics

Statistical analyses were performed using GraphPad Prism (v 10.1.0). The results are expressed as the mean ± SEM. Statistical analyses of the differences among groups were performed using two-way ANOVA for multiple comparisons with Tukey’s post-hoc test and one-way ANOVA with Tukey’s post-hoc test when appropriate (i.e. gene analysis comparison between sorted populations). Values of p < .05 were considered significant (*), p < .01 very significant (**) and p < .001 extremely significant (***).

## Declarations

### Ethics approval

All tissues or cells were obtained with written consent and under local ethic board’s approval. Use of all human materials was approved by the McGill University Health Centre Research Ethics Board, under project #2019-5374 for iPSCs.

### Consent for publication

Not applicable.

### Availability of data and materials

Not applicable.

## Authors’ contributions

VECP conceptualized the study and designed all *in vitro* experiments, interpreted data, and prepared the manuscript. AC and GJB performed some *in vitro* experiments, acquired and analyzed some *in vitro* data, and helped with manuscript preparation. JS performed all the FACS experiments and heled with manuscript preparation. ZY generated the SOX10^mO^ iPSC line. VS helped with the design of some experiments, provided the astrocyte media and method, and contributed to manuscript preparation. JPA, GB and TMD substantially contributed to the design of the study and to critical manuscript revision. All authors read and approved the final version of the manuscript.

## Acknowledgements

We thank lab members Irina Shlaifer, Wolfgang Reintsch, Andrea Krahn and Genevieve Dorval for technical or administrative assistance. We acknowledge the McGill microscopy platform for their support, Figures 1A and 4a were created with BioRender.com.

## Additional files

**Additional file 1: Supplementary figures**

Supplementary figures S1-S6.

