## Supplementary Figures for "The Use of a SOX10 Reporter Towards Ameliorating Oligodendrocyte Lineage Differentiation from Human Induced Pluripotent Stem Cells"

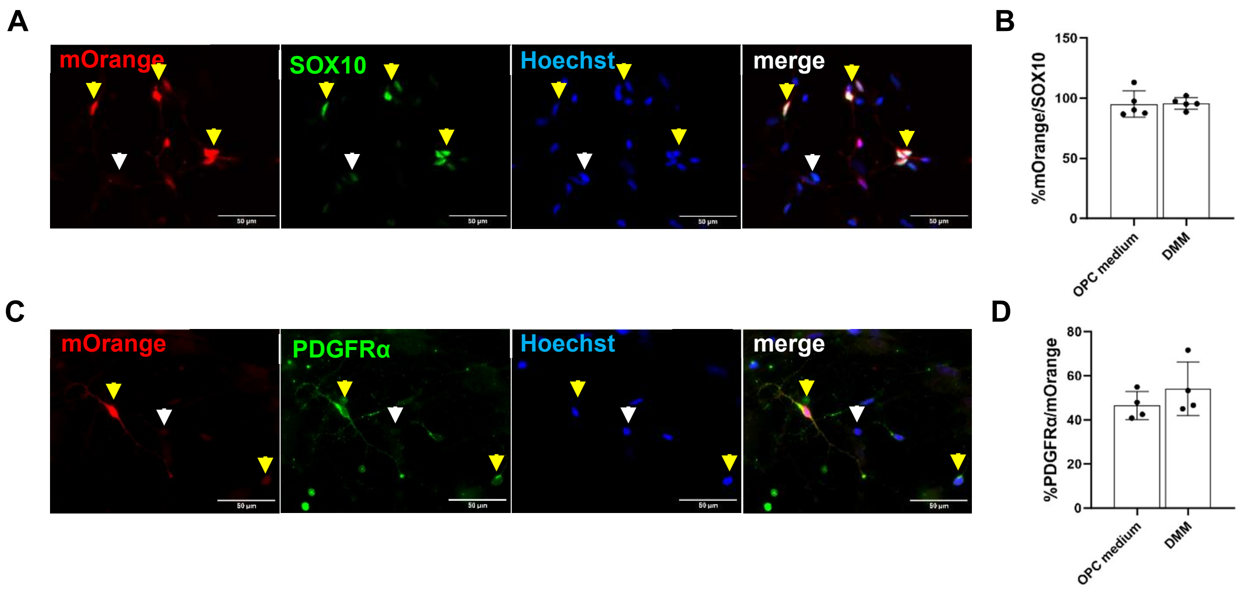


**FIGURE S1. Validation of SOX10$^{mO}$ iPSC-derived OPCs.** (A) Immunofluorescence of OPCs obtained at *DIV* 75; cells positive for mOrange are in red and SOX10 in green; the yellow arrows indicate some of the cells co-expressing mOrange and SOX10, white arrows highlight SOX10-positive cells not co-expressing mOrange; nuclei (in blue) are marked with Hoechst33342, scale bar: 50µm. (B) Histogram representing the percentage of SOX10-positive cells co-expressing mOrange in OPC medium and DMM; n=5, one-way ANOVA (ns, p=0.979, Tukey's multiple comparison). (C) Immunofluorescence of *DIV* 75 OPCs showing colocalization of mOrange, in red, with PDGFRα, in green; yellow arrows show co-localization, white arrows show mOrange-positive cells not expressing PDGFRα; scale bar: 50µm. (D) Histogram representing the percentage of mOrange-positive cells co-expressing PDGFRα; n=5, one-way ANOVA (ns, p = 0.855, Tukey's multiple comparison).

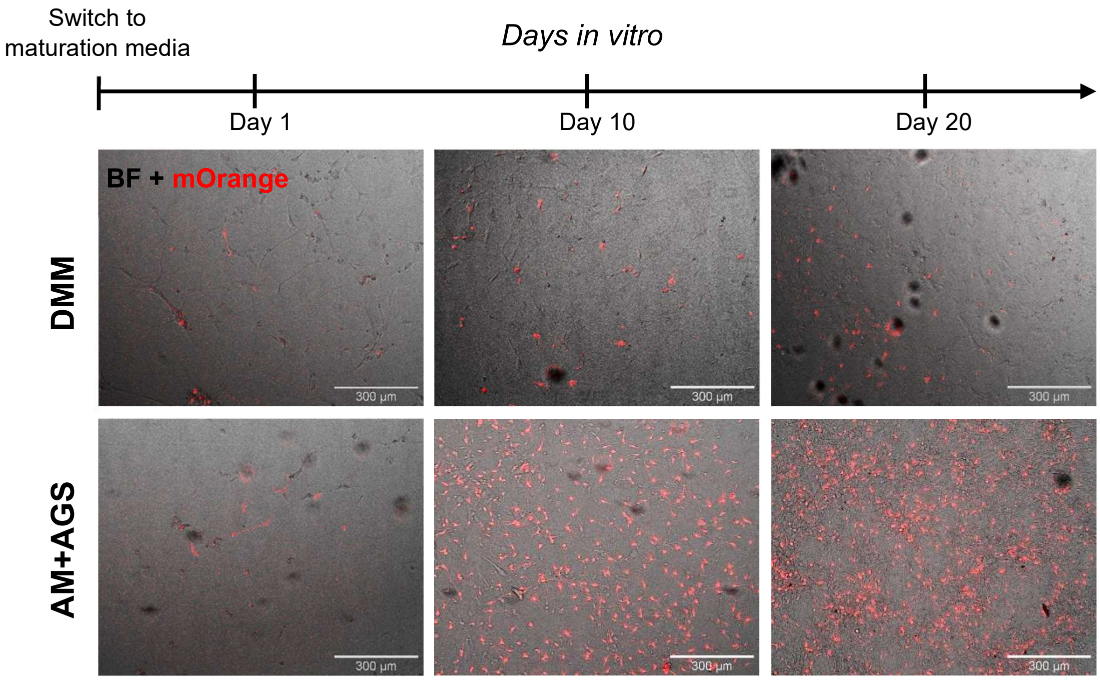


**FIGURE S2. Increase of mOrange-positive cells percentage during maturation.** Brightfield images of cells differentiated over 20 days in DMM (upper panels) and AM+AGS (lower panels) respectively. The expression of mOrange is represented in red. Scale bar: 300µm.

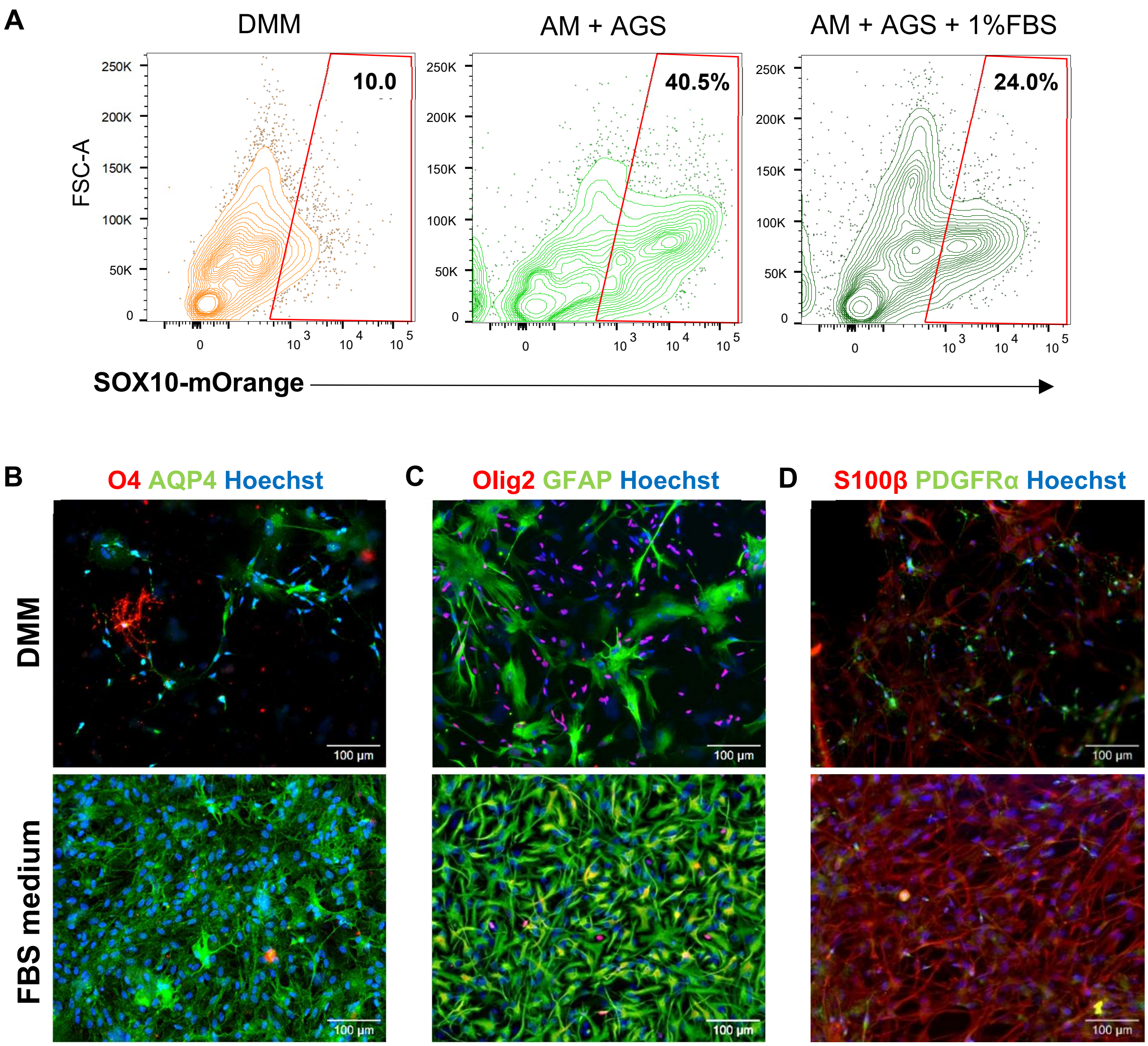


**FIGURE S3. The presence of FBS in the culture media enhances the astrocytic commitment.** (A) Flow cytometry analysis of SOX10-mOrange cells after21 days exposure to DMM, AM+AGS or FBS medium; (B-D) Immunofluorescence analysis comparing cells exposed for 21 *DIV* to DMM (upper panels) vs cells differentiated in AM+AGS supplemented with 1% FBS for the same period. (B) the astrocytic marker Aquaporin-4 (AQP4) is visualized in green and the OL marker O4 in red; (C) the astrocytic GFAP is showed in green with Olig2 in red highlighting OL nuclei; (D) S100β (in red) marks cells of the astrocytic lineage and PDGFRα (in green) marks OPCs. Nuclei are detected with Hoechst (in blue). Scale bars: 100µm.

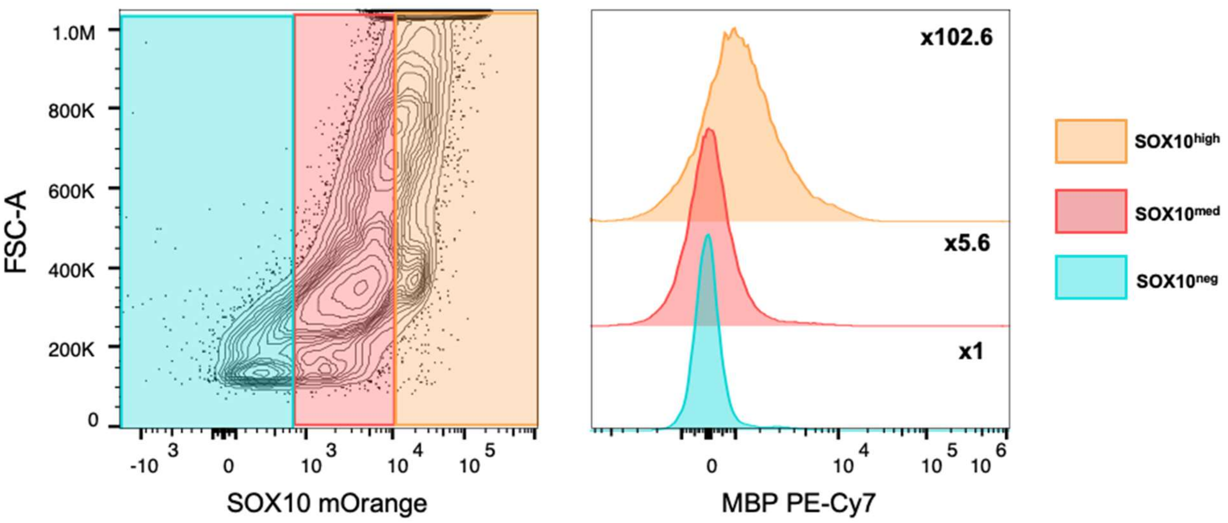


**FIGURE S4. Flow cytometry analysis of MBP expression in relation with mOrange intensity.** The left panel shows three different groups of cells based on SOX10-mOrange expression (indicated in the legend on the right); right panel shows the intensity of MBP in each of the groups. Fold-change in comparison with SOX10-negative group (SOX10$^{neg}$) is indicated next to each curve.

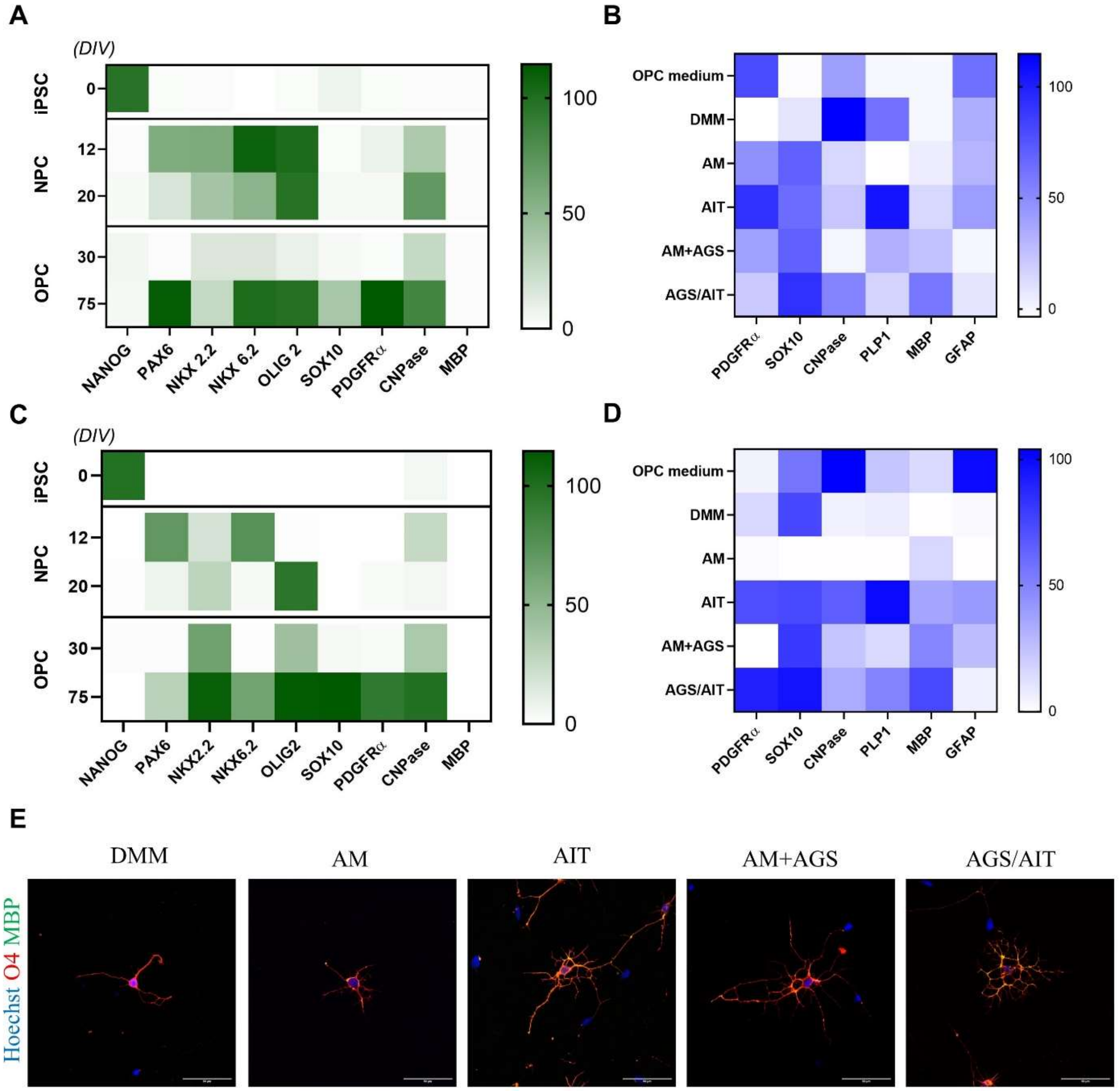


**FIGURE S5. Analysis of different maturation media in 3450 and 81280 cell lines.** (A) Heatmap of normalized gene expression in different phases of OPC induction in 3450 cell line (n=4, two-way ANOVA). The *days-in-vitro* (*DIV*) are shown on the left border; (B) normalized gene expression heatmap showing a comparison of different maturation media in 3450 cell line (n=4, two-way ANOVA); (C) Heatmap of normalized gene expression in different phases of OPC induction in 81280 cell line (n=2, two-way ANOVA). The *days-in-vitro* (*DIV*) are shown on the left border; (D) normalized gene expression heatmap showing a comparison of different maturation media in 81280 cell line (n=2, two-way ANOVA); (E) Day 95 OLs from 81280 cell line in each maturation medium staining O4 (red) and MBP (green) at 40X (scale bars: 50µm).

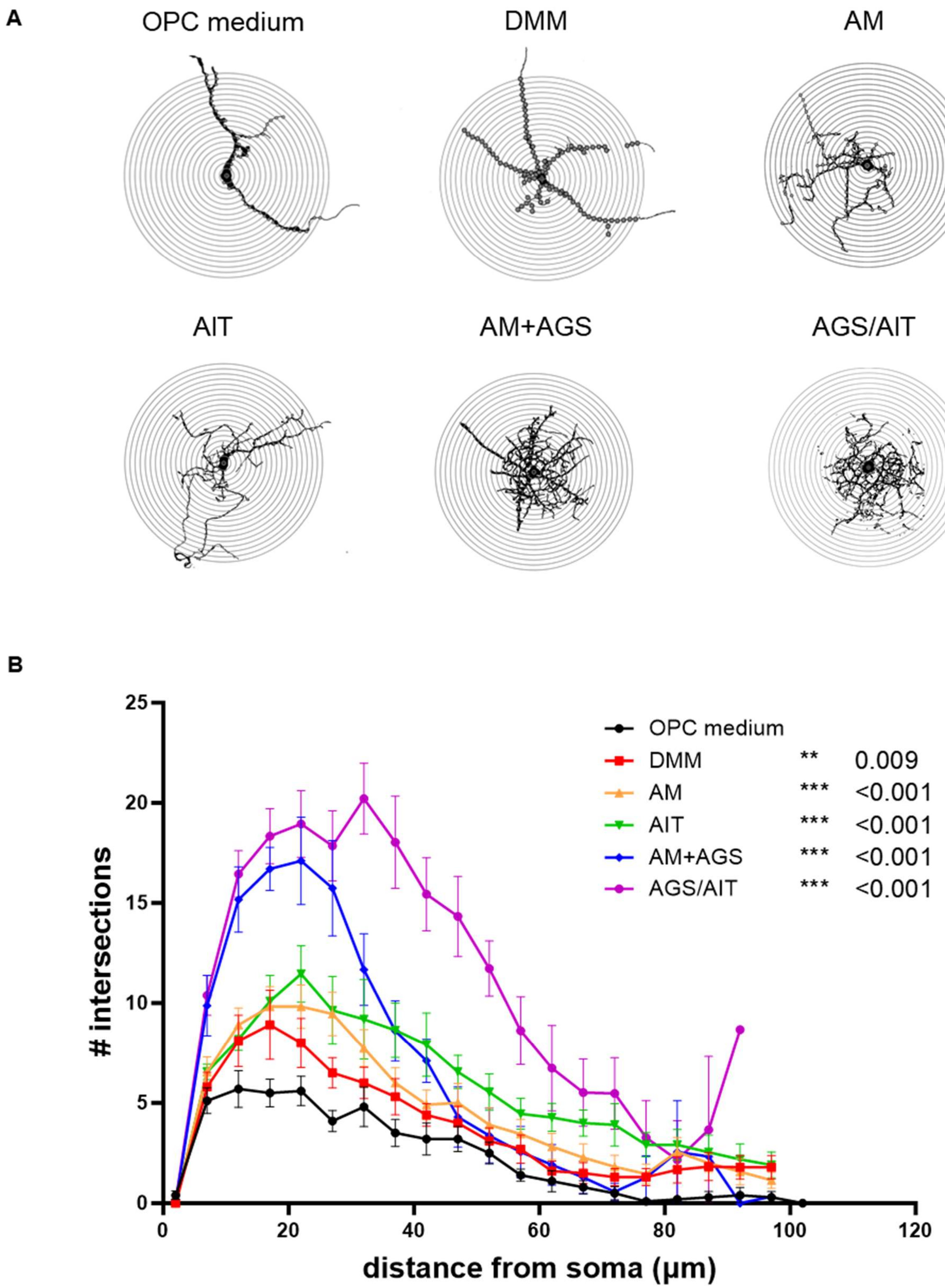


**FIGURE S6. Detailed Sholl analysis of SOX10$^{mO}$ cells in different maturation media.** (A) representative interaction maps showing the cell morphology in different maturation media; (B) graph representing the average number of intersections at each radius from the cell body in different media conditions. Significance is expressed as *p-value* of average n.intersections in comparison with OPC medium and is shown in the legend (n=10, one-way ANOVA with Tukey's multiple comparisons).
